# Probing the effect of PEG-DNA interactions and buffer viscosity on tethered DNA in shear flow

**DOI:** 10.1101/2025.04.05.647409

**Authors:** Fatema Tuz Zohra, Huda Al-Zuhairi, Jefferson Reinoza, HyeongJun Kim, Andreas Hanke

## Abstract

DNA flow-stretching is a widely employed, powerful technique for investigating the mechanisms of DNA-binding proteins involved in compacting and organizing chromosomal DNA. We combine single-molecule DNA flow-stretching experiments with Brownian dynamics simulations to study the effect of the crowding agent polyethylene glycol (PEG) in these experiments. PEG interacts with DNA by an excluded volume effect, resulting in compaction of single, free DNA molecules in PEG solutions. In addition, PEG increases the viscosity of the buffer solution. By stretching surface-tethered bacteriophage lambda DNA in a flow cell and tracking the positions of a quantum dot labeled at the free DNA end using total internal reflection fluorescence (TIRF) microscopy, we find that higher PEG concentrations result in increased end-to-end length of flow-stretched DNA and decreased fluctuations of the free DNA end. To better understand our experimental results, we perform Brownian dynamics simulations of a bead-spring chain model of flow-stretched DNA in a viscous buffer that models the excluded volume effect of PEG by an effective attractive interaction between DNA segments. We find quantitative agreement between our model and the experimental results for suitable PEG-DNA interaction parameters.

## 1. Introduction

Deoxyribonucleic acid (DNA) is a highly charged, semi-flexible polymer that stores genetic information in all living organisms. Importantly, the length of DNA is much longer than cell or nucleus sizes. For example, in each human cell, the total length of DNA confined within a nucleus of about 10 μm diameter[1] is approximately two meters [1,2]. Likewise, bacterial chromosomes are typically ∼1,000 times longer than cell sizes [3–5] The large ratio of DNA to cell dimensions requires highly dynamic and organized DNA packaging. A variety of DNA-binding proteins employ various strategies to turn long linear DNA into a dense structure to package within the cellular volume. For instance, DNA supercoiling, bending (or wrapping), and loop formation by bridging different segments of DNA are well-characterized DNA compaction mechanisms [5,6]. Additionally, some DNA-binding proteins are capable of structuring DNAs by actively extruding DNA loops in an adenosine triphosphate (ATP)-dependent manner [7–11]. It has also been shown that liquid-liquid phase separation can assist in DNA compaction [12,13]. In addition to DNA packaging, DNA compaction and decompaction play crucial roles in various aspects of cellular events, such as gene regulation by changing chromatin accessibility and protection against chemical or biochemical stress [14,15].

The conformational dynamics of polymers and biopolymers in flow are of great experimental and theoretical interest. The stretching of single, tethered DNA molecules by a uniform (nonshearing) flow was observed by fluorescence microscopy in Perkins *et al*. [16]. The dynamics of single, free (untethered) DNA molecules in steady shear flow was observed by fluorescence microscopy in Smith *et al*. [17]. The dynamics of single DNA molecules tethered to a surface in shear flow was observed experimentally by fluorescence microscopy and studied by Brownian dynamics simulations of bead-spring chains [18–20]. The study in Doyle *et al*. [18] revealed the intriguing phenomenon of large temporal fluctuations in the chain extension due to a continual recirculating motion of the chain at moderate flow strengths, referred to as *cyclic dynamics* (see also Lueth and Shaqfeh [20] and Supporting Information, Movie S1). The rheological and optical behavior of bead-rod chains in steady, linear flows was studied by Brownian dynamics simulations in Doyle *et al*. [21]. The effect of attractive surfaces on the stretching of confined tethered polymers under shear flow was studied by Brownian dynamics simulations in Ibáñez-García *et al*. [22] and reviewed in Refs. [23–25]. In the fluorescence microscopy experiments summarized above, the DNA was uniformly labeled with dye molecules, which alters the persistence length and other mechanical properties of the DNA being studied [23,24]. This problem was overcome by attaching a fluorescent quantum dot (with a diameter of a few tens nanometers) to the free end of the shear-stretched DNA and tracking the position of the quantum dot by fluorescence microscopy instead of using a DNA intercalating dye, allowing for the study of the DNA extension and the magnitude of fluctuations of the DNA as a function of the shear rate [26]. This new approach was refined by labeling DNA molecules with multiple fluorescent quantum dots at specific sites along the DNA contour and tracking their positions in time by fluorescence microscopy, revealing the dynamics of and correlations between mesoscopic subsegments of the DNA [27]. The single-molecule flow-stretching assay [28,29] has proved to be a powerful method for probing mechanisms of DNA-binding proteins and their implications in DNA [30–36] (Fig 1A) (and the related DNA curtain assay [37–39]). Another benefit of the single-molecule DNA flow-stretching assay is its high sensitivity. For example, by using this technique, we have recently demonstrated that attaching a small amino acid tag to DNA-binding proteins can alter their functional properties [40].

**FIG 1.**
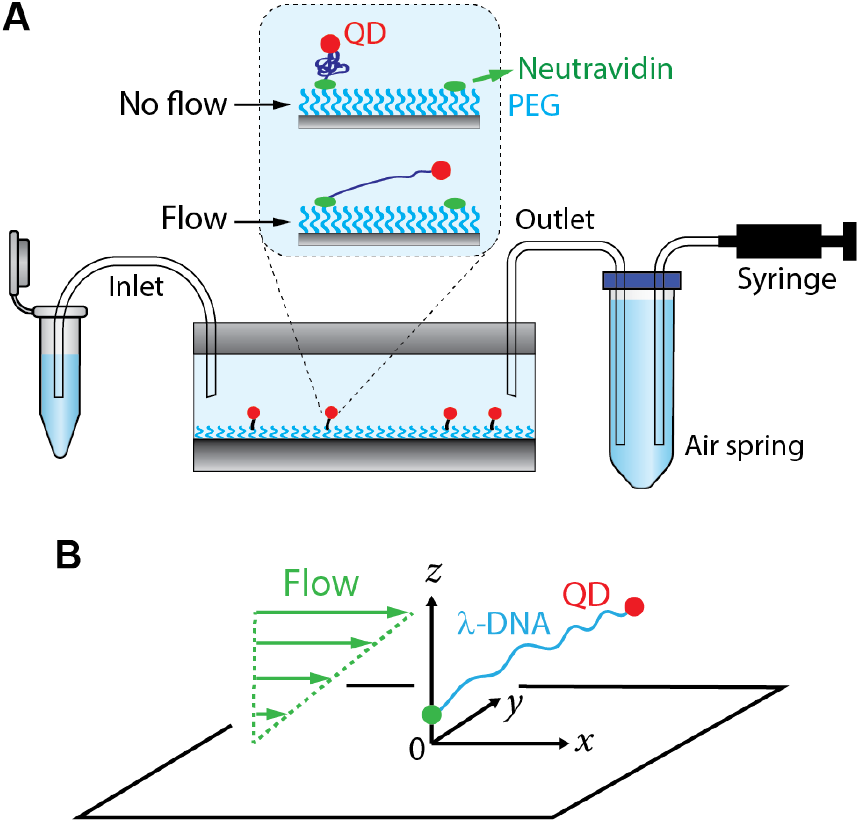
Schematics of our experimental setup and model. (A) Biotinylated quantum dot (QD)-labeled *λ*-DNAs are tethered to a surface-passivated microfluidic flow cell via neutravidin-biotin interactions. As syringe plunger is withdrawn, buffer flows from the inlet tube side to the syringe through the microfluidic flow cell. The hydrodynamic drag force leads to DNA stretching. Polyethylene glycol (PEG) on the surface minimizes unwanted nonspecific DNA and QD binding to the surface. Air spring helps maintain the flow rate. (B) Coordinate system used in our model. *λ*-DNA (blue) is tethered close to the surface at one end (green dot) and labeled by a quantum dot (QD, red) at the other end. The surface of the flow cell is in the *xy*-plane at *z* = 0. The DNA is subject to a flow field whose speed in *x*-direction increases linearly with the distance *z* from the surface (green).

Cellular environments of all living organisms are densely crowded with macromolecules including proteins, metabolites, and small solutes [41]. For instance, the concentrations of cytoplasmic macromolecules were measured to be up to 450 g/L [42,43]. Such a crowded medium resembles a thick molecular soup where the average distances between macromolecules are significantly smaller than their sizes [44]. Although proteins are typically studied *in vitro* in diluted aqueous buffers, efforts have been made to conduct experiments in crowded conditions [31,45,46]. In this study, we investigate the effect of the commonly used crowding agent polyethylene glycol (PEG) on flow-stretched DNA. PEG affects biomolecular solutions through excluded volume effects[47,48] or co-condensation [49,50], and its physical properties are well-documented [41,51,52]. In particular, free DNA molecules in solution collapse upon the addition of PEG because the thermodynamically unfavorable contact between DNA and PEG decreases the available free space for coil DNA, leading to an effective, PEG-induced attraction between DNA segments [53,54]. This effect is similar to the hydrophobic interaction of biomolecules in an aqueous solution, in which nonpolar (hydrophobic) residues seek to minimize the surface area exposed to water resulting in an effective, nonspecific attraction between hydrophobic residues. In addition to crowding, PEG also increases the solvent viscosity [41], which leads to an increased drag on flow-stretched DNA. Therefore, the influence of PEG on solvent viscosity must be taken into account in single-molecule DNA flow-stretching assays in the presence of PEG.

We examine the dynamics of single bacteriophage *λ*-DNA molecules in a microfluidic flow cell for different concentrations of PEG by labeling the free DNA end with a fluorescent quantum dot (QD) and dynamically tracking the motion of the QD in real time using total internal reflection fluorescence (TIRF) microscopy. The single-molecule DNA flow-stretching experiments show that supplementing the imaging buffer with PEG increases DNA stretching under laminar flow and reduces fluctuations of the untethered free DNA ends. The observed dynamics results from the dual role of PEG, generating both an effective attraction between DNA segments and increasing the buffer viscosity. To better understand our experimental results, we perform Brownian dynamics simulations of a bead-spring chain model of flow-stretched DNA in a viscous buffer that incorporates the PEG-induced effective attraction between DNA segments. We find quantitative agreement between our model and the experimental results for the chain extension and the strength of the fluctuations of the free DNA end for suitable PEG-DNA interaction parameters, providing proof of principle of understanding the dynamics of DNA interacting with an agent in a crowded environment.

## 2. Experimental materials and methods

### 2.1 Viscosity measurements for buffers containing PEG

Buffer viscosities were measured using a HAAKE MARS rheometer (Thermo Scientific, Waltham, MA) with a parallel plate-plate geometry, where the radii of the rotor and the lower plates were 17.5 mm and 18.0 mm, respectively. The axial gap between the plates was 1 mm. For each PEG concentration, viscosities were measured at 80 different shear rates between 0.01 (s^-1^) and 1000 (s^-1^); however, for the geometry we used in our measurements, the data obtained were not reliable at low shear rates (between 0.01-0.1 s^-1^). The measurements were repeated four or five times, and values averaged over these measurements were used. All experiments were performed at 23 °C.

The composition of the imaging buffer used in the viscosity measurements was 10 mM Tris, pH 8.0, 150 mM NaCl, and 10 mM MgCl2. Experiments were performed without PEG and with 3%, 5%, or 10% of PEG supplemented to the buffer. The Pearson correlation coefficient was calculated using Excel software.

### 2.2 DNA substrate preparation

The complementary single-strand 5’ overhangs of *λ*-phage DNA were utilized to tag one DNA end with biotin and the other with digoxigenin as described previously [32]. Biotin allows us to tether the DNA to the microfluidic flow cell surface via neutravidin-biotin interactions. The digoxigenin molecule was used to label the free DNA end with an anti-digoxigenin antibody-conjugated quantum dot 605 (Invitrogen, Waltham, MA). A biotinylated oligo was annealed to one of the complementary single-stranded overhangs and ligated, followed by annealing and ligation of a digoxigenin-oligo. Unreacted excess short oligos were removed by electrophoresis, and the DNA substrates in EB buffer (10 mM Tris, pH 8.5) were obtained by ethanol precipitation.

### 2.3 Microfluidic flow cell preparation

To minimize nonspecific DNA binding, cover glass surfaces were passivated by (3-aminopropyl)triethoxysilane (Millipore Sigma A3648, St. Louis, MO) followed by a mixture of PEG (MPEG-SVA-5000-1g) and its biotinylated version (Biotin-PEG-SVA-5000-100mg) (Laysan Bio, Arab, AL) as described previously [30–32,40]. A microfluidic flow cell was constructed by applying double-sided tapes (Grace Bio-Labs, Bend, OR) between a PEGylated cover glass and a quartz plate (Technical Glass Product, Paineville, OH). The parallel-attached double-sided tapes form a channel of defined height and width (see Section 2.4). The PE60 inlet (13 cm in length) tube attached to one end of the channel was dipped into a tube containing a buffer. The outlet tube attached to the other end of the channel was connected to the syringe on a syringe pump (Harvard Apparatus, Holliston, MA) through an air spring (Fig 1A).

### 2.4 Single-molecule DNA flow-stretching assay

The height of the channel of a microfluidic flow cell, *h* = 0.12 mm (*z*-direction in Fig 1B), was determined by the thickness of the tape. The channel width, *w* = 1.8 mm (*y*-direction in Fig 1B, was the same as in our previous studies [30,31,40]. One end of 48.5-kb *λ*-phage DNA [55] was tethered to the sample chamber via neutravidin-biotin interactions, and the other end was tagged with a fluorescence quantum dot. The tethered DNAs were stretched by applying laminar flow at 50 μL/min generated by a syringe pump.

The flow speed in the flow direction (*x*-direction in Fig 1B) at height *z* from the surface of the flow cell is given by a parabola [56], i.e.,

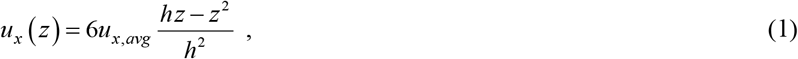

where 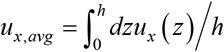 is the average flow speed and *u x* (0) = *u*_*x*_ (*h*) = 0 . In our experiments, we used a flow rate of *Q* = 0.833 *μ*L/s in a flow cell of cross-sectional area *w* · *h* = 0.216 mm^2^ corresponding to an average flow speed of *u*_*x*,*avg*_ = *Q* (*w* · *h*) = 3.856 mm/s.

### 2.5 Single-molecule DNA flow-stretching and data analysis

A small (∼4%) percentage of the PEGylated surface of the microfluidic flow cell contained biotin molecules. Addition of 0.25 mg/mL neutravidin followed by quantum dot-labeled *λ*-DNA resulted in tethering of the DNA onto the surface at the biotin-tagged ends. Unlabeled quantum dots and untethered DNAs were washed away by flowing imaging buffer (10 mM Tris, pH 8.0, 150 mM NaCl, and 10 mM MgCl2). When there was no buffer flow for at least two minutes, the movie acquisition was initiated. The average quantum dot position corresponds to the DNA tether point in the absence of flow. Subsequently, we stretched the DNAs by turning on the imaging buffer flow (flow rate *Q* = 0.833 *μ*L/s) (Supporting Information, Movie S2). The imaging buffer without polyethylene glycol (PEG) was switched to the buffer supplemented with a given amount (3% and 5%) of PEG while the flow rate remained constant. All experiments were performed on the IX-83 total internal reflection fluorescence (TIRF) microscope (Evident Scientific, Olympus, Waltham, MA) equipped with a 532 nm laser (Coherent, Santa Clara, CA). Micro-manager software [57] was employed to record images, and regions-of-interest (ROI) of DNAs were set using FIJI software [58]. Although the size of the quantum dots is only about a few tens of nanometers, their EMCCD images (point spread functions, PSFs) are a few hundred nanometers due to diffraction. Thompson *et al*. formulated an equation that shows how accurately the position of a fluorophore can be determined [59]. Later, Yildiz *et al*. demonstrated that, if enough photons are collected, fluorophore positions can be determined even with nanometer accuracy after fitting the PSF with a two-dimensional Gaussian function [60]. We used custom-written MATLAB software based on one-dimensional Gaussian fitting along the DNA length to determine the quantum dot positions. The MATLAB codes are available in our previous publication [30].

### 2.6 Averages over ensembles of single DNA molecules

Because of the heterogeneous nature of DNA stretching and fluctuations, the procedure described in Section 2.5 was repeated for 49 individual DNA molecules for each of the cases where the buffer without PEG was switched to buffers containing 3% and 5% PEG, respectively. This resulted in data for *N* = 98 DNA molecules for the buffer without PEG and *N* = 49 DNA molecules for buffers containing 3% and 5% PEG, respectively. Measured values for the mean extensions ⟨*xi*⟩ and standard deviations Δ*xi* for DNA molecules *i* = 1,…, *N* were averaged according to

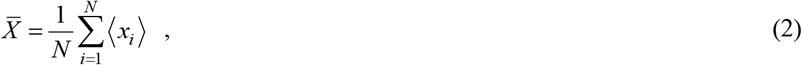

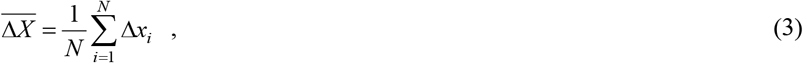

where the overbar symbol indicates averages over the ensemble of *N* molecules. The changes of 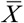 and 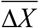 upon adding to the buffer without PEG a given amount (3% or 5%) of PEG were statistically evaluated through the nonparametric Mann-Whitney test (Wilcoxon rank-sum test) using Prism software (GraphPad, San Diego, CA).

## 3. DNA model and computational methods

### 3.1 Bead-spring model for DNA in a flow cell

To better understand the effect of the PEG-induced effective attraction between DNA segments and the buffer viscosity in our DNA flow-stretching experiments, we performed Brownian dynamics simulations of a bead-spring chain with parameters corresponding to a nearly inextensible worm-like chain such as DNA. The experimental setup (Fig 1A) is modeled as shown in Fig 1B. A linear DNA chain is modeled by a bead-spring model consisting of *N* + 1 beads centered at positions 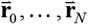 connected by *N* stiff harmonic springs with equilibrium length *a* and spring constant *k*_*s*_ (the bead radius does not enter our analysis) (Fig 2). As in our experiments, the chain is attached at one end close to the surface of the flow cell (bead 0 at position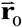) while the other end is moving freely (bead *N* at position 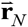). In our Brownian dynamics simulations (detailed below), we use chains with *N* = 50 beads, corresponding to *a* = *L N* = 330 nm for the contour length *L* = 16.49 *μ*m of the *λ*-DNA used in our experiments, and we assume that the chain is attached at 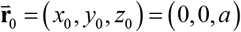.The spring constant *k*_*s*_ is chosen larger than all other force constants to account for the fact that biopolymers like DNA are nearly inextensible [61]. As a result, stretching modes relax fast compared to other dynamic processes, so that the simulated chain essentially behaves like an inextensible polymer chain. On the other hand, *k*_*s*_ cannot be chosen too large as larger *k*_*s*_ require smaller discretization time steps Δ*t* to ensure numerical stability of the Brownian dynamics simulations, thus limiting the maximal simulation time *t*_*max*_ . As a compromise, we chose the value *k*_*s*_ = 5000 *k*_*B*_*T* / *a*^2^ where *T* is the temperature and *k*_*B*_ is the Boltzmann constant.

**FIG 2.**
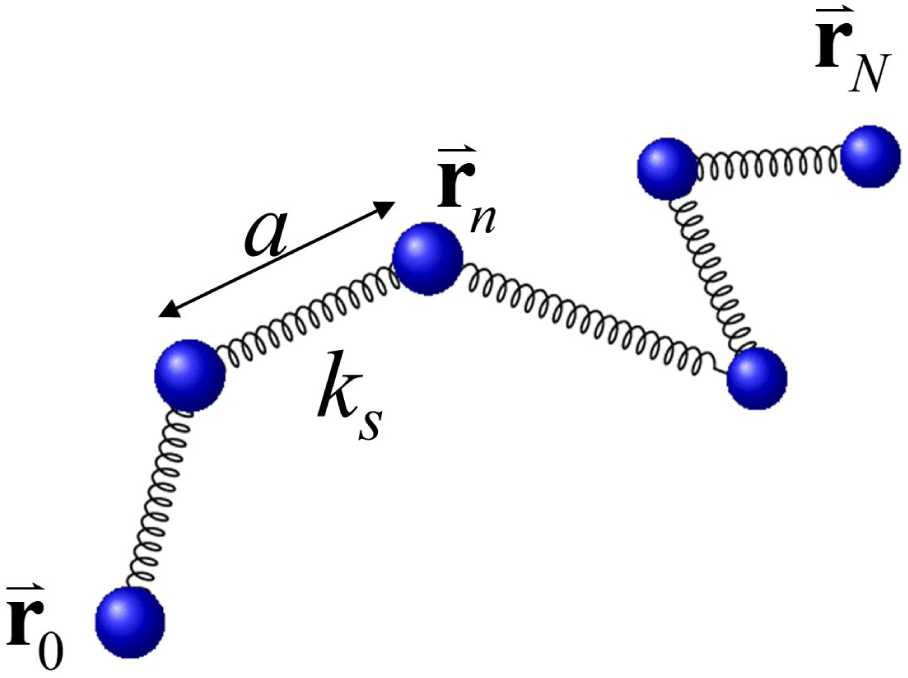
Bead-spring model for linear DNA. *N* + 1 beads are connected by springs with spring constant *k*_*s*_ and equilibrium length *a* . The chain is tethered at 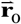 close to the surface of the flow cell and 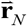 is the position vector of the free end. (see Fig 1B).

The non-hydrodynamic forces on the beads of the model chain include (i) the spring forces, (ii) the external force by the surface to which the chain is tethered, and (iii) the forces generated by the PEG-induced attractive interaction between DNA segments. The interaction of the model chain with the viscous solvent is described by (iv) a drag force and (v) a Brownian force due to random collisions of the solvent with the beads. The forces (i) – (v) are detailed below.

(i) Spring forces: The total elastic (el) force on a bead *n* within the chain is given by

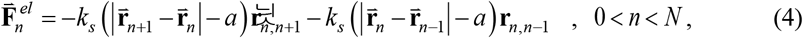

where *k*_*s*_ is the spring constant and 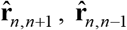 are unit vectors from bead *n* +1 and *n* −1 to bead *n* , respectively. The elastic force on bead *N* at the free end of the chain is given by 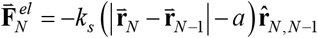 . As discussed above, *a* = 330 nm for the *λ*-DNA used in our experiments modeled by a chain with 50 beads, and we used the value *k*_*s*_ = 5000 *k*_*B*_*T* / *a*^2^ in our simulations.

(ii) Surface interaction: For the repulsive interaction between the polymer chain and the surface (s) we assume a truncated soft-wall potential of the form

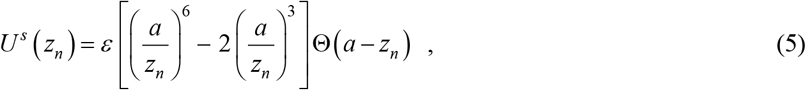

Where *z*_*n*_ is the distance of bead *n* from the surface (Fig 1B), and Θ(*a* − *zn*) = 1 for 0 ≤ *z*_*n*_ ≤ *a* and Θ(*a* − *z*_*n*_) = 0 for *z*_*n*_ > *a* . In our Brownian dynamics simulations, we set *ε* = *k*_*B*_*T* . The force 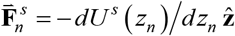 on bead *n* resulting from Eq. (5) is given by

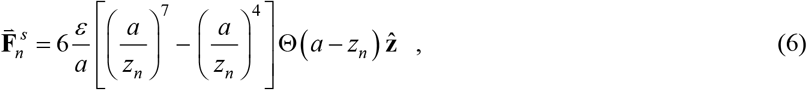

where 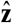 is a unit vector perpendicular to the surface pointing into the flow cell. 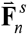 is repulsive for 0 ≤ *zn* < *a* , equal to zero for *z*_*n*_ ≥ *a* , and continuous at *z*_*n*_ = *a* . According to the size of our flow cell and the length of the *λ*-DNA used in our experiments, the DNA only interacts with the surface at *z* = 0 to which it is tethered (Fig 1B).

(iii) PEG-induced effective attraction between DNA segments: Since there is no generally accepted energy potential for the PEG-induced effective attraction between DNA segments, we use a phenomenological energy potential between DNA segments that captures the main effect (compare, e.g., Refs. [62,63]). Accordingly, for our bead-spring model of DNA, we assume a pairwise attractive interaction between beads *n* and *n* ‘ described by the potential energy

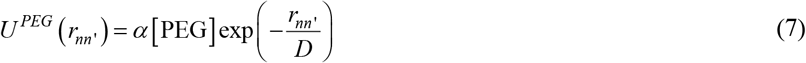

where 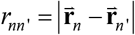 is the distance between the beads located at 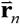 and 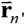 respectively, *α* is a constant, [PEG] is the PEG concentration in percent (%), and *D* is the exponential decay length of the PEG-induced attractive interaction between DNA segments. Equation (7) incorporates the assumptions that the PEG-induced attraction between DNA segments (corresponding to the beads in our model) is proportional to [PEG] and decays exponentially with the bead separation. In a model with atomistic resolution, the decay length *D* is expected to be related to the radius of gyration (size) *R*_*g*_ of the PEG polymers, which in our experiment is a few nanometers [64,65]; however, in our coarse-grained DNA model using a bond length of *a* = 330 nm (Fig 2) the decay length *D* should be comparable to the bond length *a* to obtain a notable PEG-induced change in the mean DNA length *x*_*e*_ . We performed Brownian dynamics simulations for two different values of the decay length, namely *D* = *a* and *D* = *a* 2 , to show that the constant *α* in Eq. (7) is indeed independent of [PEG] and the buffer viscosity *η* , regardless of the value of *D* (more precisely, *α* depends on *D* but for given *D* is independent of [PEG] and *η* ). Using Eq. (7) the net force 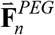 on bead *n* due to the PEG-induced attraction between nearby beads is given by

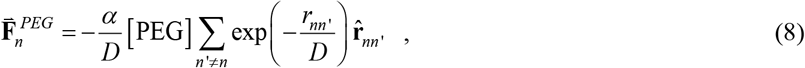

where the sum includes all beads *n*′ different from *n* and 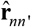 is the unit vector directed from bead *n*′ to *n* . For the elongated conformations occurring in our simulations of flow-stretched DNA, only beads *n*′ close to bead *n* along the contour of the chain contribute significantly to the net force in Eq. (8). To reduce the computational cost in our simulations, we therefore restricted the sum in Eq. (8) to beads *n*′ with 1 ≤ *n* − *n*′ ≤ 5 ; including more than 5 nearest-neighboring beads did not change our results within the statistical error of our simulations.

(iv) Viscous drag: The drag force between the viscous solvent and the polymer chain is represented by Stoke’s drag force on the beads of the model chain. The drag force on the bead *n* is given by

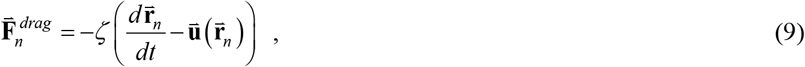

where *ζ* is the drag coefficient, 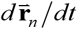 is the velocity vector of bead *n* , and 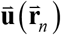 is the unperturbed flow velocity vector of the solvent at position 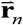 of bead *n* . Assuming laminar shear flow in *x*-direction (Fig 1B), the velocity vector of the solvent at position 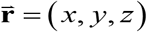 in the flow cell has the form

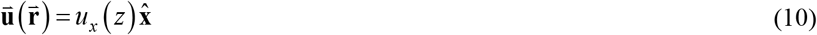

where 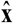 is a unit vector in *x*-direction and the velocity profile *u*_*x*_ ( *z*) is given by Eq. (1). The shear rate of the fluid is given by

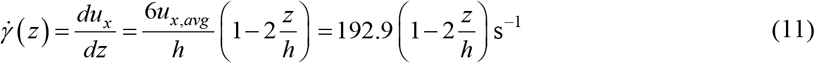

where we used *u*_*x*,*avg*_ = 3.856 mm/s and *h* = 0.12 mm for our flow cell (see text below Eq. (1)). In our DNA flow stretching experiments the ratio ⟨ *z*_*e*_*/ h*⟩ , where ⟨ *z*_*e*_⟩ is the average height of the free end of the DNA chain above the surface (Fig 1B), is always smaller than 0.002; thus, the term 2 *z /h* in Eq. (11) is only a small correction to the leading term. In our Brownian dynamics simulations, it was therefore justified to neglect this correction, and we assumed a constant shear rate

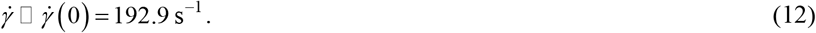

Accordingly, in Eq. (9) we used the approximation

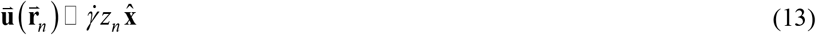

with the constant shear rate 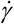 in Eq. (12).

### 3.3 Langevin equation and Brownian dynamics simulations

The total force on bead *n* of the model chain (Fig 2) includes the forces 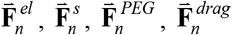 described in (i) – (iv) above, and (v) a Brownian (thermal) force 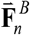 due to random collisions of the solvent particles with the bead. According to Newton’s second law of motion and neglecting inertia, the net force 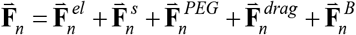 on any bead *n* is zero, resulting in the Langevin equation for bead *n* [66,67]

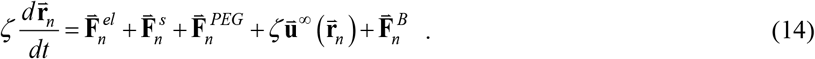

The Brownian force 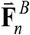 is a stochastic (random) force, taken from a Gaussian distribution in our simulations. For the dynamics to satisfy the fluctuation-dissipation theorem [66,67], the expectation values of the Brownian force with components ν = *x, y, z* are given by

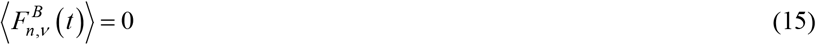

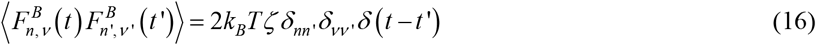

with the drag coefficient *ζ* in Eq. (9); *Δ* _*nn*’_ is the Kronecker delta, *Δ* (*t* − *t*′) is the Dirac delta function, *T* is the temperature, and *k*_*B*_ is the Boltzmann constant.

For the numerical integration of Eq. (14) we use an Euler-Maruyama iteration scheme by discretizing time in small steps Δ*t* [68]. We write Eq. (14) in terms of reduced (unitless) variables by expressing lengths in units of *a* , energies in units of *k*_*B*_*T* , forces in units of *k*_*B*_*T* / *a* , and times in terms of the unit time

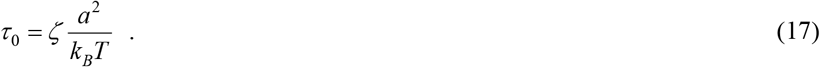

*τ* _0_ is the time for a single, free bead with drag coefficient *ζ* to diffuse a distance *a* in a solvent of temperature *T* . Discretizing time in steps Δ*t* the delta function in Eq. (16) becomes *Δ* (*t* − *t*′) = 1 Δ*t* for *t* = *t*′ and zero otherwise. Each time step *i → i* + 1 advances the position 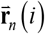 of bead *n* at time *t* (*i*) = *i*Δ*t* according to

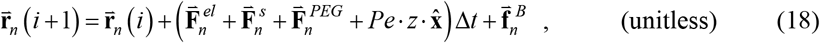

where *Pe* is the Péclet number (see next paragraph below). According to Eqs. (15) and (16), the stochastic impulse 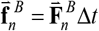 is determined by the correlations

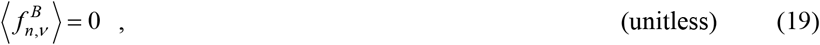

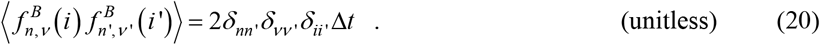

To simplify notation in Eqs. (18)–(20) we use the same symbols for the reduced (unitless) variables defined above as for their dimensionful counterparts, as explicitly indicated; otherwise, we will use a tilde symbol to indicate reduced variables (see next paragraph below). Equation (20) implies that the stochastic impulse is independent of all beads *n* , spatial directions *v* = *x, y, z* , and time steps *i → i* +1 . The strength of the Brownian random force is characterized by the variance 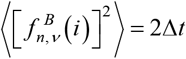 (unitless variables). In our numerical implementation, we used unitless time steps Δ*t* = 10^−4^ in Eq. (18), and values 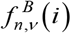 were drawn from a Gaussian distribution with zero mean and variance 2Δ*t* independently for each *n* , *v* , and *i* .

### 3.3 Relaxation time and Weissenberg number

In Eq. (18) arises the unitless Péclet number

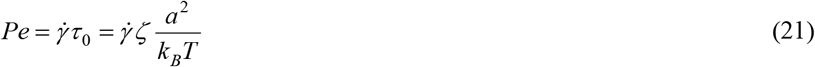

corresponding to the ratio of the time *τ* _0_ in Eq. (17) for a single bead to freely diffuse a distance *a* to the time scale 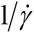 set by the shear rate 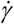 of the flow in Eq. (12). For a long polymer chain rather than a single bead, a more relevant parameter is the unitless Weissenberg number

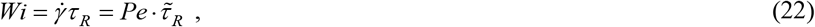

where *τ* _*R*_ is the longest relaxation time of the chain and 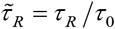; in Eq. (22) and below, we indicate reduced (unitless) variables by a tilde symbol to distinguish them from their dimensionful counterparts. In our simulations, we determined 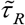 in two different ways (Fig 3A and 3B): (A) By the exponential decay of the autocorrelation function 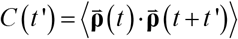 at equilibrium (i.e., without flow and at thermal equilibrium), where 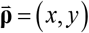 is the two-dimensional projection of the three-dimensional end-to-end vector 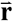 on the *xy* - plane (Fig 1B). Fitting *C* (*t*′) to a single exponential 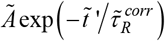, where 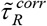 and Ã are unitless fit parameters and 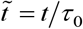 , yields 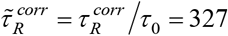 (with Ã = 30.57) (Fig 3A) (we used the autocorrelation function of 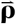 instead of 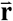 because 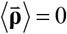 by symmetry, allowing for a fit of *C* (*t*′) to a single exponential with only two free parameters). (B) By the exponential decay of the mean value 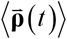 *to* equilibrium starting from a non-equilibrium state at *t* = 0 ; in our simulations, the initial non-equilibrium state of the chain consisted in a fully extended conformation along the *x*-direction at *t* = 0 , allowing the chain to relax to equilibrium for *t* > 0 (in the absence of flow). Fitting 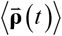 to a single exponential 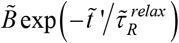 , where 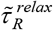 and 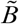 are unitless fit parameters, yields 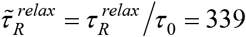 (with 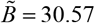) (Fig 3B). According to Onsager’s regression hypothesis (or, more generally, the fluctuation-dissipation theorem) [66,67], we expect 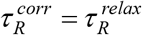,which holds for our simulation results with an error of about 3%. To proceed, in this paper we use the average of the values obtained by the two methods, i.e.,

**FIG 3.**
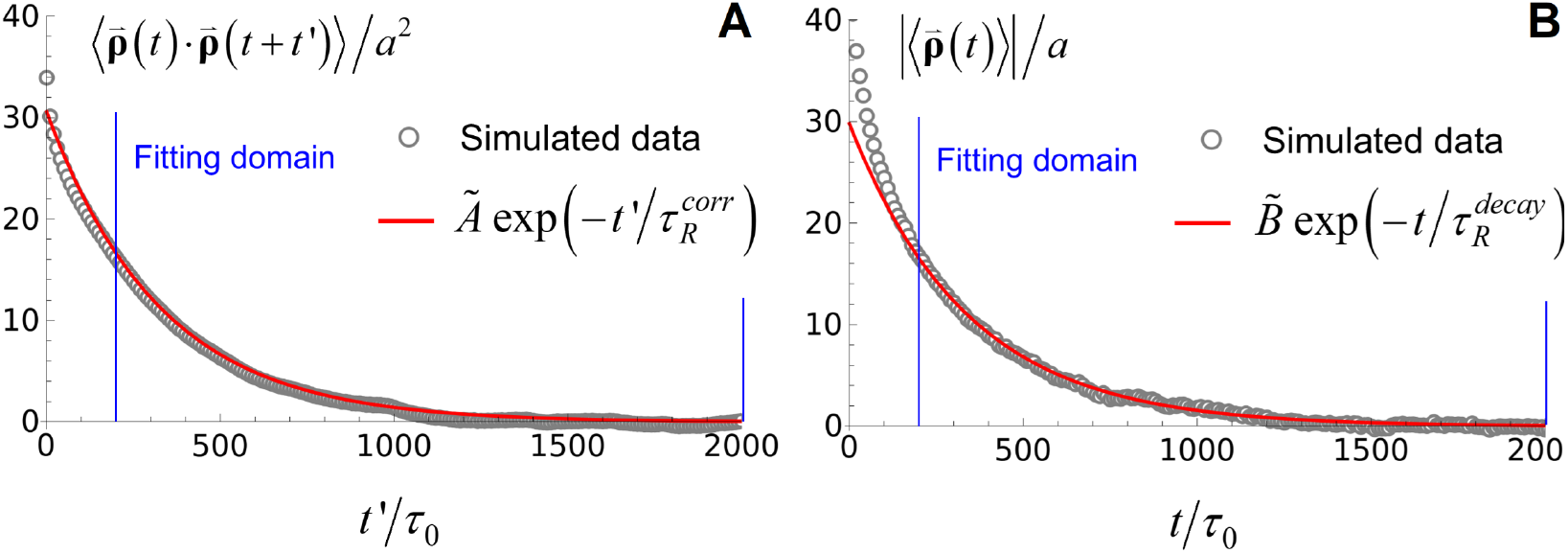
Determination of the longest relaxation time of the polymer chain, *τ* _*R*_ , in our simulations. *τ* _*R*_ is determined by (a) the autocorrelation function 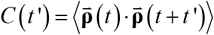 *at* equilibrium and (b) the relaxation of 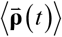 *to* equilibrium, where 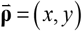 is the projection of the chain end on the *xy* – plane [Fig 1(b)]. *C* (*t*′) and 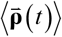 were fitted by least squares to single exponentials in the domain 200 ≤ *t*′ *τ* _0_ , *t τ* _0_ ≤ 2000 (vertical blue lines), yielding 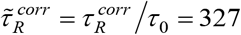 and 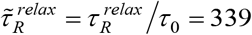 , respectively.

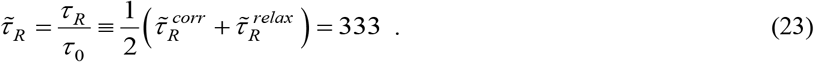

## 4. Results

### 4.1 PEG leads to higher buffer viscosities

To confirm that adding PEG to a buffer solution increases the viscosity, we first measured buffer viscosities while changing the shear rate using a rheometer with a parallel plate-plate configuration. In the rotational rheometer, one plate rotates relative to the other, inducing shear on the fluid between the two plates.

In our experiments, for the imaging buffer (see Section 2) without PEG, the viscosity η remained roughly constant (0.914 ± 0.02 cP) for shear rates 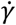 in the range 60–500 s−1 (Supporting Information, Figures S1A and S1B), suggesting that the buffer behaves as a Newtonian fluid. Supplementing the buffer with 3%, 5%, or 10% concentrations of PEG, respectively, increases η in this range of shear rates as follows: 1.675 ± 0.05 cP (3% PEG), 2.396 ± 0.03 cP (5% PEG), and 5.680 ± 0.04 cP (10% PEG) (Supporting Information, Fig S1C), using the notation (mean value) ± (standard deviation, SD) (Pearson’s correlation coefficient: 0.976), and these PEG-containing buffers are again Newtonian fluids (see Table 1). The viscosity measurements directly demonstrate that PEG increases the viscosity of a buffer, which therefore needs to be taken into account in our single-molecule DNA flow-stretching assay.

**Table 1.** Measured buffer viscosity and DNA length as a function of PEG concentration.

| [PEG] | $\langle\eta\rangle \pm \text{SD}$ (cP) | $\bar{X} \pm \text{SD}$ (SEM) ($\mu\text{m}$) | $\overline{\Delta X} \pm \text{SD}$ (SEM) ($\mu\text{m}$) |
| --- | --- | --- | --- |
| 0% | $0.914 \pm 0.02$ | $11.897 \pm 1.249$ (0.126) | $0.3966 \pm 0.1095$ (0.0111) |
| 3% | $1.675 \pm 0.05$ | $12.446 \pm 1.545$ (0.221) | $0.3538 \pm 0.1118$ (0.0160) |
| 5% | $2.396 \pm 0.03$ | $12.554 \pm 0.819$ (0.117) | $0.2652 \pm 0.0601$ (0.0086) |
| 10% | $5.680 \pm 0.04$ | no data | no data |

### 4.2 Experimental DNA flow-stretching at different PEG concentrations

We experimentally investigated to what extent the end-to-end length of flow-stretched DNA depends on the PEG concentration. One end of *λ*-phage DNA was tethered to the surface of the flow cell and the other end was labeled with a fluorescent quantum dot (QD) (Fig 1A).

Measured buffer viscosity ⟨*η* ⟩, average end-to-end DNA length 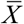 in *x*-direction (Eq. (2)), and standard deviation 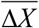 of the DNA end position in *x*-direction [Eq. (3)], as a function of PEG concentration in our DNA flow-stretching experiments. Errors are given by the standard deviation (SD) followed by the standard error of the mean (SEM) in parentheses as indicated. SD and SEM were determined from *N* = 49 independent measurements for 3% and 5% [PEG] and *N* = 98 independent measurements for 0% [PEG] (Fig 4 and 5, and Section 2).

**FIG 4.**
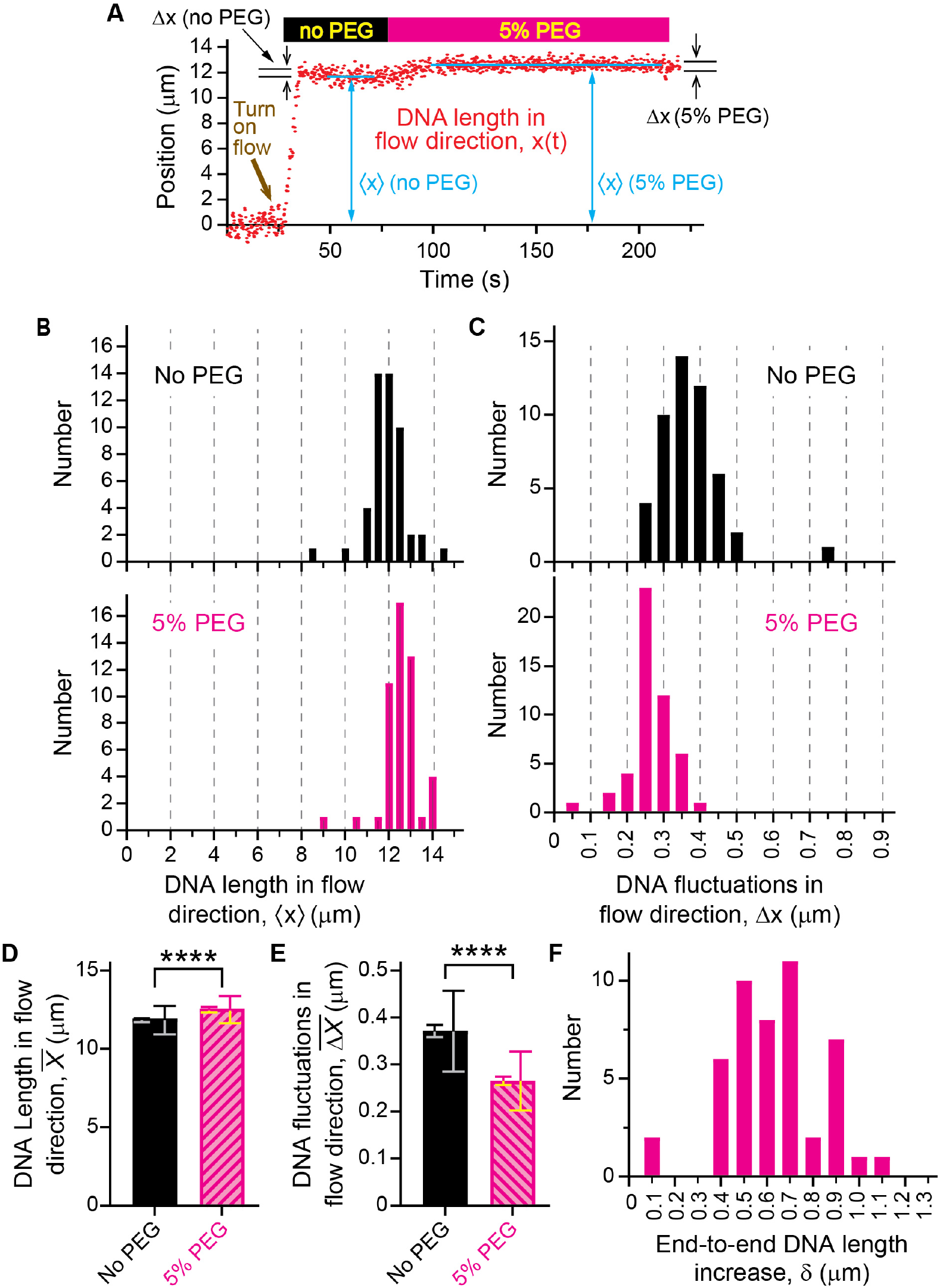
Supplementing 5% PEG increases the end-to-end flow-stretched DNA length and reduces fluctuations. (A) Representative graph of quantum dot (QD) position versus time for an individual DNA molecule. Data collection of QD positions *x* (red dots) vs time began in the absence of flow and continued while turning on flow in the *x*-direction leading to DNA stretching. Data were collected for an imaging buffer without PEG and for a buffer supplemented with 5% PEG. For a given PEG concentration, the mean value ⟨*x*⟩ (horizontal blue lines) and the standard deviation (SD) Δ*x* were determined from the corresponding set of QD positions. (B) Histograms of mean values ⟨*x*⟩ from independent measurements on 49 DNA molecules for buffers without PEG (upper panel) and 5% PEG (lower panel). (C) Same as (B) for SDs Δ*x*. (D) Mean value 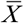 of mean values ⟨*x*⟩ for the ensemble of 49 DNA molecules corresponding to the histograms in (B) for buffers without PEG and 5% PEG, respectively (see Eq. (2)). The error bars show the SDs (larger error bars) and the standard errors of the mean (SEMs, smaller error bars) corresponding to the histograms in (B). ****: *p* < 0.0001 by Mann-Whiteney test. (E) Same as (D) for the mean value 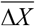 of SDs Δ*x* corresponding to the histograms in (C) (see Eq. (3)). (F) Histogram of the relative increase in chain extensions *δ* = ⟨*x*⟩ (5%) − ⟨*x*⟩ (0%) obtained for the 49 individual DNA molecules upon supplementing the buffer with 5% PEG.

**FIG 5.**
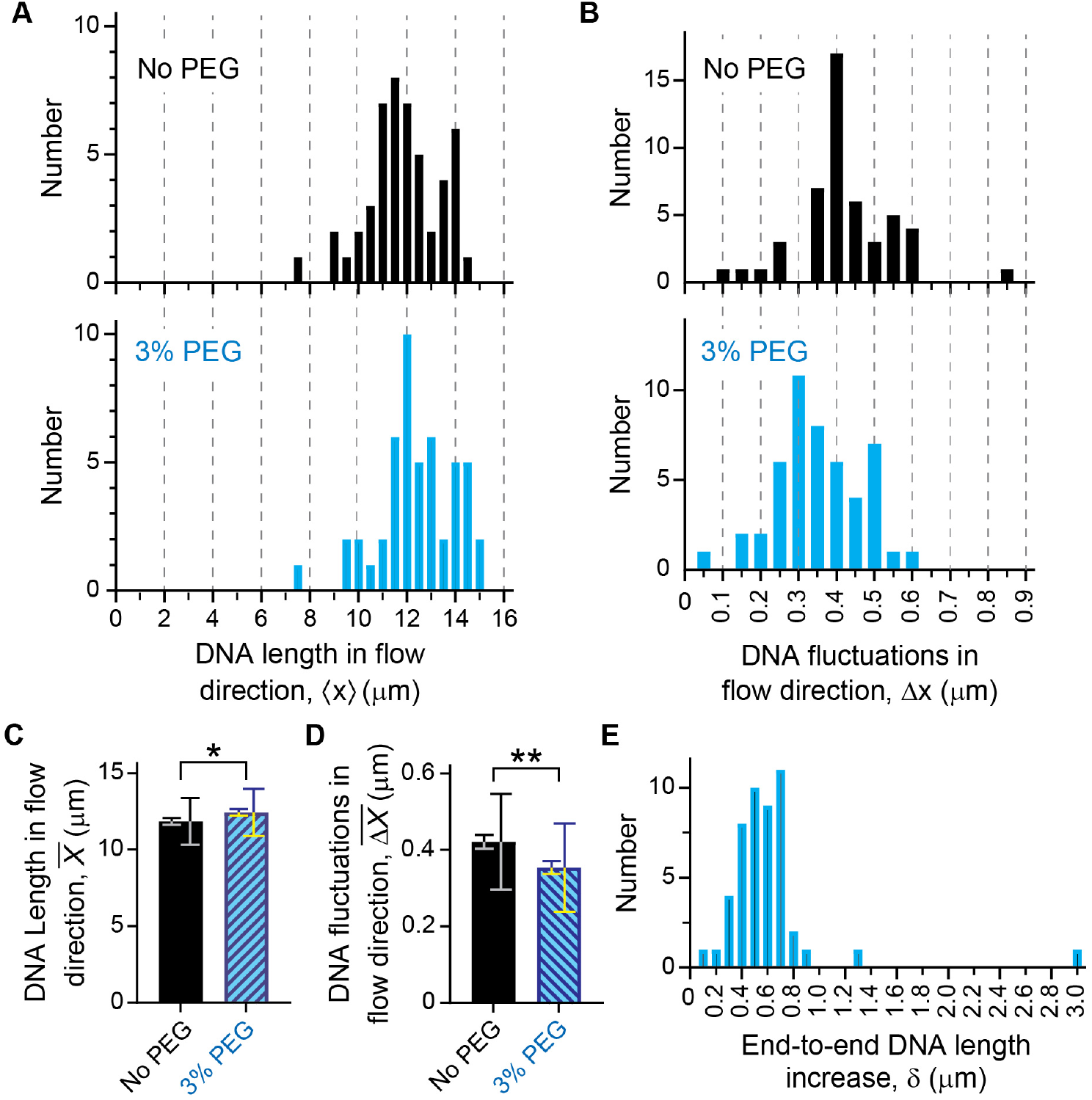
Supplementing the buffer with 3% PEG increases the end-to-end flow-stretched DNA length and reduces fluctuations. (A) Histograms of mean values ⟨*x*⟩ from independent measurements on 49 DNA molecules for buffers without PEG (upper panel) and 3% PEG (lower panel). (B) Same as (A) for the standard deviations (SDs) Δ*x*. (C) Mean value 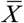 of mean values ⟨*x*⟩ for the ensemble of 49 DNA molecules corresponding to the histograms in (A) for buffers without PEG and 3% PEG, respectively (see Eq. (2)). The error bars show the SDs (larger error bars) and the standard errors of the mean (SEMs, smaller error bars) corresponding to the histograms in (A). *: 0.01 < *p* < 0.05 by Mann-Whiteney test. (D) Same as (C) for the mean value Δ*X* of SDs Δ*x* corresponding to the histograms in (B) (see Eq. 3). **: 0.001 < *p* < 0.01 by Mann-Whiteney test. (E) Histogram of the relative increase in chain extensions *δ* = ⟨*x*⟩ (3%) − ⟨*x*⟩ (0%) obtained for the 49 individual DNA molecules upon supplementing the buffer with 3% PEG.

For a given DNA molecule and in the absence of flow, the average QD position in *x*-direction, ⟨*x*⟩, was approximately that of the DNA tether point. Applying flow in *x*-direction stretches the DNA (Supporting Information, Movie S2), and the resulting increase of the average end-to-end DNA length ⟨*x*⟩ was obtained by comparing average QD positions before and after the DNA was stretched by flow (Fig 4A, red dots). We started with an imaging buffer without PEG followed by a buffer supplemented with 5% PEG (Fig 4A). We also calculated the standard deviation Δ*x* (fluctuations) along the *x*-direction of the QD positions on the flow-stretched DNA. The average end-to-end length ⟨*x*⟩ and fluctuations Δ*x* were obtained for both buffer conditions (no PEG and 5% PEG). Although multiple DNA molecules are observed in a single field of view, each of them behaves slightly differently in terms of stretching and fluctuations. Observing individual DNA molecules allows the detection of heterogeneous events. Thus, these measurements were performed for 49 different DNA molecules. Figures 4B and 4C show histograms of the resulting mean chain extensions ⟨*xi*⟩ and standard deviations Δ*xi* , respectively, for no PEG and 5% PEG. The histograms in Fig 4B are broader than the fluctuations for a single DNA molecule in Fig 4A. This is because errors in determining DNA tether point positions, in addition to fluctuations of QD positions, affect the DNA end-to-end length measurements (see Section 2). Figures 4D and 4E show the averages 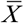 and 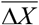 , respectively, over these values (see Eqs. (2) and (3)), again for no PEG and 5% PEG. These data show that the addition of 5% PEG increases the average chain extension 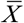 (Fig 4D) and reduces the average fluctuations 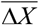 (Fig 4E) compared to the buffer without PEG in a statistically significant way (*p*-value < 0.0001 obtained by Mann-Whitney test). Finally, Fig 4F shows a histogram of the relative increase in chain extensions, *Δ*_*i*_ = *x*_*i*_ (5%) − *x*_*i*_ (0%) obtained for the individual DNA molecules upon supplementing the buffer with 5% PEG, resulting in an average 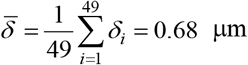 . Similar results were obtained from experiments comparing DNA length and fluctuations for buffers without PEG and with 3% PEG (Fig 5A-5E).

### 4.3 The dual role of PEG in DNA flow-stretching experiments

As discussed in Section 1, PEG plays a dual role in DNA flow-stretching experiments: On the one hand, PEG tends to reduce the DNA length by inducing an effective, attractive interaction between DNA segments; on the other hand, PEG increases the DNA length by increasing the buffer viscosity *η* . To distinguish these two counteracting effects, we assume that, in general, the mean DNA length ⟨*x*⟩ in the flow direction (Fig 1B) is a function of *η* and the PEG concentration, which are considered as independent variables:

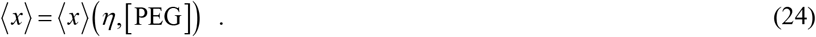

That is, the dependence of ⟨*x*⟩ on the buffer viscosity is described by the first argument *η* , whereas the dependence of ⟨*x*⟩ on the PEG-induced attractive interaction between DNA segments is described by the second argument [PEG] . For example, ⟨*x*⟩ (*η* ,0) corresponds to the *η* - dependence of ⟨*x*⟩ in the absence of an attractive interaction between DNA segments. In our DNA flow-stretching experiments (indicated by the subscript “exp” below), the buffer viscosity *η* is determined by the PEG concentration, i.e., *η*_exp_ = *η* ([PEG]) (Table 1, second column); thus, the measured mean DNA length 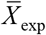 for the ensemble of 49 DNA molecules [see Eq. (2)] is effectively a function of [PEG] only (Table 1, third column), which by Eq. (24) can be written as

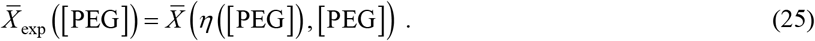

Measurements for bacteriophage lambda DNA indicate that if the electrolyte concentration is high enough (∼10 mM of NaCl and above), the relaxation time *τ* _*R*_ in Eq. (22) is proportional to the viscosity *η* of the solvent [66,69], i.e., *τ* _*R*_ = *cη* , where *c* is a constant. The buffer used in our single-molecule flow-stretching assay meets this condition (10 mM Tris pH 8.0, 150 mM NaCl, 10 mM MgCl2, 0.2 mg/mL bovine serum albumin, see Section 2). Assuming that the relaxation time of the PEG is much faster than *τ* _*R*_ so that *τ* _*R*_ is determined by the viscosity, the Weissenberg number in Eq. (22) is given by

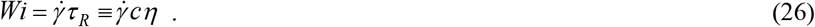

Since the shear rate 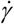 in our experiments is also constant [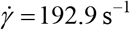, see Eq. (12)], we obtain

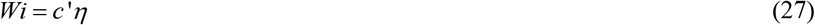

Where 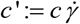 is another constant. Equations (25)–(27) allow for a comparison of our experimental results for the mean DNA length, 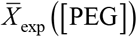 in Eq. (25), with the Brownian dynamics simulations for a single chain as follows (Fig 6A and 6B). First, we simulated the fractional extension *ξ* = *x L* as a function of *Wi* in the absence of PEG, i.e., *α* = 0 in Eq. (8) (using the Péclet number 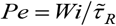 in Eq. (18) with 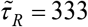 from Eq. (23)), where *L* is the contour length of the chain. This yields the simulated (sim) function *ξ*_sim_ (*Wi*) (Fig 6A), labels on bottom axis). Using the condition *ξ*_sim_ (*Wi*_0_) ≡ *ξ*_exp_ (*η*_0_) , where Eq. (18) with 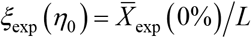 is the mean fractional extension measured for a flowing buffer without PEG and *η*_0_ = 0.914 cP (first line in Table 1), yields the Weissenberg number associated with the viscosity *η*_0_ , and thereby the constant *c*′ in Eq. (27),

**FIG 6.**
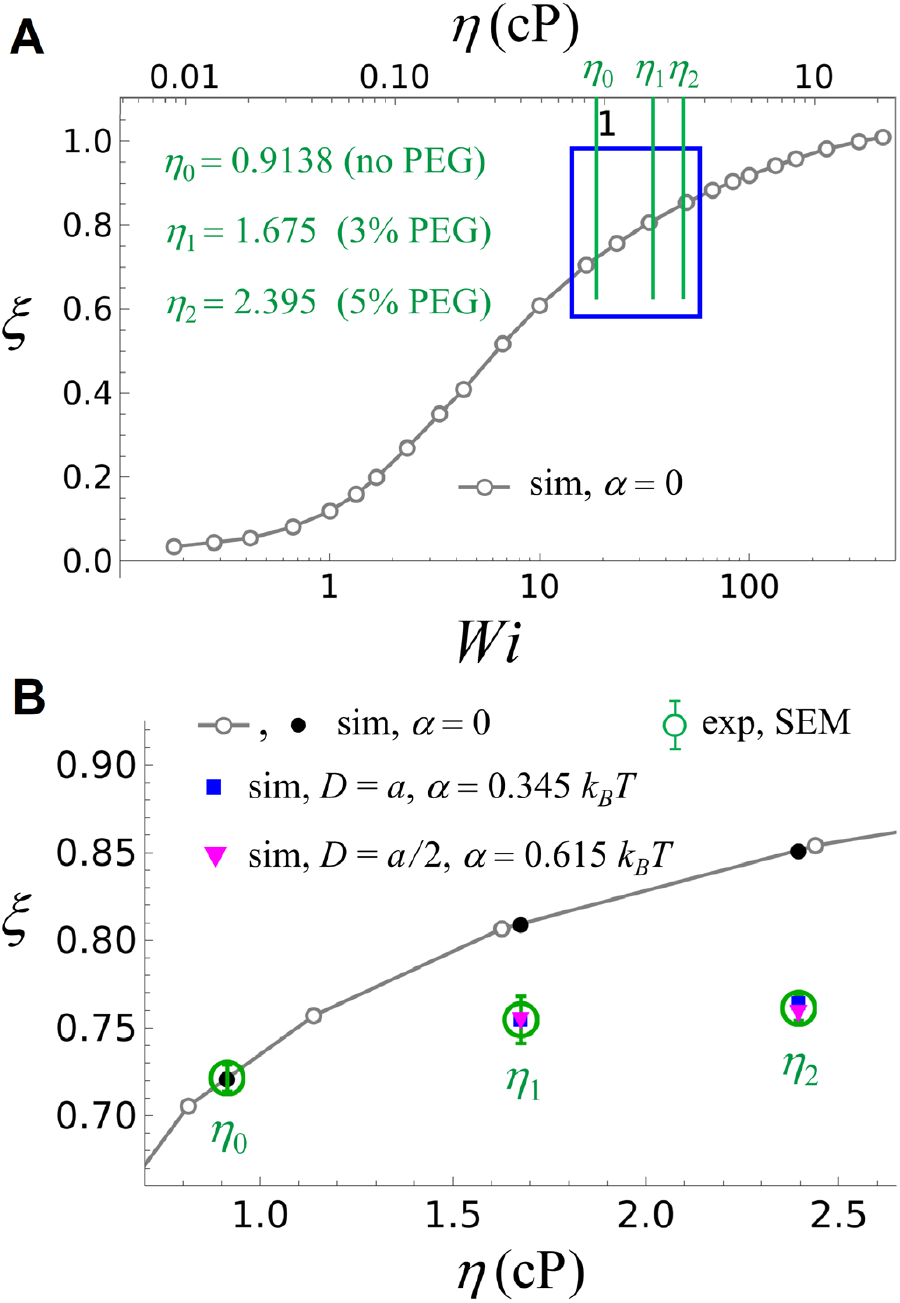
Fractional extension of flow-stretched DNA. Comparison of the fractional extension *ξ* obtained in the simulations (sim, *ξ* = ⟨*x*⟩ / *L*) and the experiments [exp, 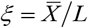,see Eq. (2) and Figures 4, 5]. (A) Simulated *ξ* versus *Wi* (bottom axis) and *η* = *Wi c*′ (top axis) [where *c*′ = 20.48 cP^−1^ , Eq. (28)] in the absence of a PEG-induced attractive interaction between DNA segments (*α* = 0 in Eq. (8)). *η*-values for buffers with 0%, 3%, and 5% PEG used in the experiments (Table 1) are indicated by green vertical lines labeled *η*_0_ ,*η*_1_,*η*_2_ , respectively. (B) Magnification of the *η*-region 0.7cP ≤ *η* ≤ 2.6 cP highlighted by the blue box in (A). Experimental *ξ*-values for 0%, 3%, and 5% PEG are indicated by green circles with error bars (SEM, Table 1). The black dots show simulated *ξ*-values for *η*_0_ ,*η*_1_,*η*_2_ in the absence of a PEG-induced attractive interaction between DNA segments. Blue and magenta symbols are simulated *ξ*-values for 3% PEG (*η*_1_) and 5% PEG (*η*_2_) using ( *D* = *a* , *α* = 0·345 *k*_*B*_*T*) and ( *D* = *a*/2*α* =0·615 *k*_*B*_*T*) in Eq. (8), respectively.

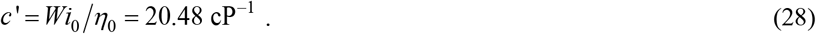

From *ξ*_sim_ (*Wi*) and using Eqs. (27) and (28), we obtain the simulated function *ξ*_sim_ (*η*) = ⟨*x*⟩ (*η*,0) *L* , i.e., the simulated fractional chain extension as a function of *η* in the absence of PEG (see Eq. (24)) (Fig 6A, labels on top axis). Thus, in Fig 6B, the agreement of *ξ* (*η*_0_) between the simulated value (leftmost black dot) and the measured value (green circle with error bars) results from the condition *ξ*_sim_ (*Wi*_0_) = *ξ*_exp_ (*η*_0_) used to determine the constant *c*′ in Eq. (28).

Next, we considered a buffer with 3% PEG corresponding to the measured viscosity value *η*_1_ = 1.675 cP (second line in Table 1). According to Eqs. (7) and (8), the PEG-induced attractive interaction between DNA segments is proportional to the PEG concentration and a constant *α* . The latter depends on the exponential decay length *D* in Eqs. (7) and (8), but for given *D* is independent of [PEG] and *η*. To find *α* for given *D* (where *D* = *a* and *D* = *a /*2 in our simulations) we used the condition

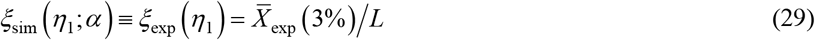

where *ξ*sim (*η*1;*α*) is the simulated fractional chain extension for *η*_1_ = 1.675 cP and given values for *α* and *D* in Eq. (8), and *ξ*_exp_ (*η*_1_) is the measured fractional chain extension for 3% PEG (see Table 1 and Eq. (25)). Equation (29) yields *α* = 0.345 *k*_*B*_*T* for *D* = *a* (Fig 6B, blue symbol for *η*_1_) and *α* = 0.615 *k*_*B*_*T* for *D* = *a/* 2 (Fig 6B, magenta symbol for *η*_1_). Thus, in Fig 6B, the agreement of *ξ* (*η*1 )between the simulated values (blue and magenta symbols) and the measured value (green circle with error bars) results from the condition in Eq. (29) used to determine *α* for given *D* .

Finally, we considered a buffer with 5% PEG corresponding to the measured viscosity value *η*_2_ = 2.396 cP (third line in Table 1). Since *α* for given *D* was determined by Eq. (29) the simulated values *ξ* (*η*_2_) for *D* = *a* (Fig 6B, blue symbol for *η*_2_) and *D* = *a/* 2 (Fig 6B, magenta symbol for *η*_2_) were obtained without any adjustable parameters. The agreement with the corresponding experimental results (green circle with error bars) provides evidence for the validity of our DNA model (Eqs. (14)–(16)) incorporating the PEG-induced attractive interaction between DNA segments by Eqs. (7) and (8).

Figure 7 shows a comparison between the standard deviation Δ*x* of the DNA end (Fig 1B) along the *x*-direction obtained in the simulations for a single chain (sim) and the average 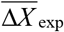 in Eq. (3) for the ensemble of 49 DNA molecules in the experiments (Fig 4 and 5). The meaning of the symbols is the same as in Fig 6. According to polymer theory, the ratio *ϕ* = Δ*x R*_*e*_ , where

**FIG 7.**
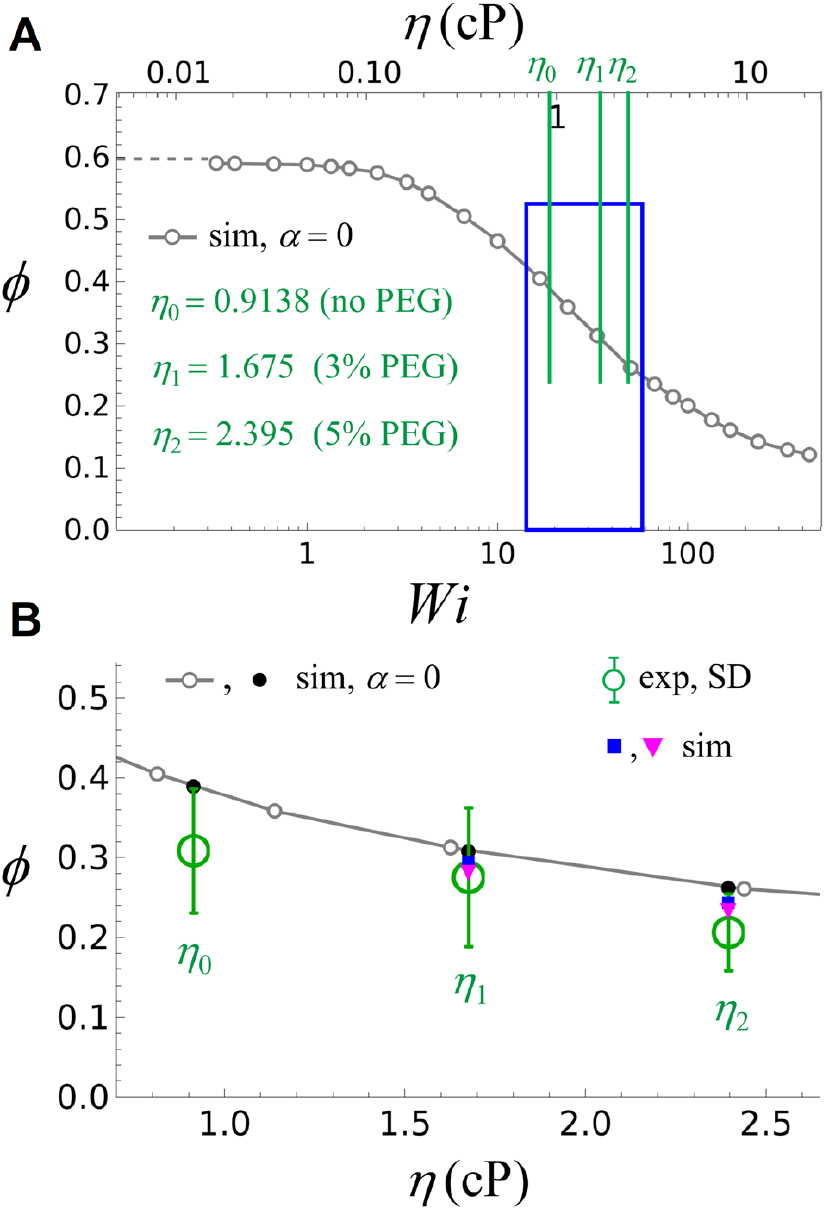
Fluctuations of the free DNA end of flow-stretched DNA. Comparison of the standard deviation (fluctuations) of the DNA end (Fig 1B) along the *x*-direction obtained in the simulations (sim, *ϕ* = Δ*x R*_*e*_) and the experiments [exp, 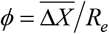,see Eq. (3) and Figures 4, 5] where *R*_*e*_ is the root mean square end-to-end distance of the chain. The meaning of the symbols is the same as in Fig 6. (A) Simulated *ϕ* vs *Wi* (bottom axis) and *η* (top axis) in the absence of a PEG-induced attractive interaction between DNA segments. (B) Magnification of the *η*-region 0.7cP ≤ *η* ≤ 2.6 cP highlighted by the blue box in (A). Experimental *ξ* -values for 0%, 3%, and 5% PEG are indicated by green circles with error bars (SD, Table 1). The black dots show simulated *ξ* -values for *η*_0_ ,*η*_1_,*η*_2_ in the absence of a PEG-induced attractive interaction between DNA segments. Blue and magenta symbols are simulated *ξ* -values for 3% PEG (*η*_1_) and 5% PEG (*η*_2_) using the same values for *α*, *D* as in Fig 6 (i.e., without adjustable parameters).

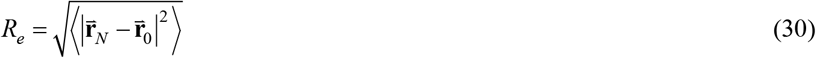

is the root mean square end-to-end distance of the free chain at equilibrium (i.e., without the surface of the flow cell and without flow, see Fig 2), is a universal function of *Wi* [18,20]. Figure 7A shows the simulated ratio *ϕ* = Δ*x R*_*e*_ vs *Wi* (bottom axis) and η (top axis) in the absence of a PEG-induced attractive interaction between DNA segments. Here, 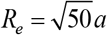 for our model chain with *N* = 50 segments of length *a* (Fig 2). Figure 7B is a magnification of the *η*-region 0.7cP ≤ *η* ≤ 2.6 cP highlighted by the blue box in Fig 7A. Experimental *ϕ* **-**values for 0%, 3%, and 5% PEG are indicated by green circles with error bars (standard deviation). These values were determined as follows. Experimental values for Δ*x* are given in Table 1. To find *R*_*e*_ for the *λ*-DNA used in our experiments, we used 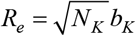 where *N*_*K*_ is the number of Kuhn segments (K) and *b*_*K*_ is the Kuhn length of the DNA. Using the DNA persistence length *ℓ* _*P*_ = 50 nm corresponding to the buffer in our experiments (see Section 2) and the contour length *L* = 16.49 *μ*m of the *λ*-DNA gives *b*_*K*_ = 2*ℓ* _*P*_ = 0.1*μ*m, thus *N*_*K*_ = *L/ b*_*K*_ = 165 and *R*_*e*_ = 1.284 *μ*m. The blue and magenta symbols in Fig 7B are simulated *ϕ* **-**values for 3% PEG (*η*_1_) and 5% PEG (*η*_2_) using the same values for **α** , *D* as determined for Fig 6, i.e., without adjustable parameters. Both the experimental and the simulated results show that Δ*x* (fluctuations of the DNA end position, see Fig 1B) decreases with increasing PEG concentration. The simulated results for *ϕ* = Δ*x /R*_*e*_ agree with the experiments within the standard error, providing further evidence for the validity of our DNA model incorporating the PEG-induced attractive interaction between DNA segments by Eqs. (7) and (8). However, for Δ*x* (Fig 7) the influence of the attractive interaction between DNA segments is less pronounced than for the chain extension ⟨*x*⟩ (Fig 6).

## 5. Discussion

We studied the dynamics of singly-tethered flow-stretched *λ*-DNA molecules by supplementing different concentrations of PEG to one of the commonly used single-molecule buffer conditions (10 mM Tris, pH 8.0, 150 mM NaCl, and 10 mM MgCl2). For this purpose, we labeled the free DNA end with a fluorescent quantum dot (QD) and dynamically tracked the motion of the QD in real time using fluorescence microscopy. Out of different crowding reagents commonly used in *in vitro* experiments, we were particularly interested in the effects of PEG due to its twofold properties: (1) While another commonly used crowding agent, sucrose, shows no evidence of DNA compaction [70], PEG can compact DNA [71] by inducing an excluded volume effect, generating an effective attraction between DNA segments [53,54]; (2) PEG increases the buffer viscosity, which can increase DNA stretching under flow. The high sensitivity of our single-molecule DNA flow-stretching assay [28,29] allowed us to resolve these counteracting effects of PEG on DNA experimentally. Furthermore, our Brownian dynamics simulations of a bead-spring chain model of flow-stretched DNA in a viscous buffer that incorporates the PEG-induced attractive interaction between DNA segments results in quantitative agreement with the experimental results for suitable PEG-DNA interaction parameters. When a PEG-induced attractive interaction between DNA segments was not considered (*α* = 0 ; Fig 6), deviations of *ξ*-values between experiments and simulations were prominent. This result underscores the importance of considering both counteracting effects of PEG. In summary, our approach provides proof of principle of understanding the dynamics of DNA interacting with an agent in a crowded environment. This is relevant for probing mechanisms of DNA-binding proteins and their implications in DNA [30–36] because the cellular environment in which these proteins act is crowded.

Our approach can be refined by labeling DNA molecules with multiple fluorescent quantum dots at specific sites along the DNA contour [27] and by using a higher-resolution model for the DNA in the Brownian dynamics simulations such as oxDNA [72]. Furthermore, our approach combining singly-tethered DNA stretching experiments with Brownian dynamics simulations lays the groundwork for future DNA stretching research. For example, in single-molecule DNA-protein studies, highly negatively charged casein and bovine serum albumin (BSA) are commonly supplemented to a buffer to minimize nonspecific adsorption of proteins onto the microfluidic flow cell [73,74]. Building on previous reports of the effects of negatively charged proteins and colloids on DNA compaction [75,76], our approach is expected to provide a deeper understanding of the effects of those unbound charged reagents in solution to the flow-stretched DNA compaction by a DNA-binding protein. Results obtained by exploring different flow cell channel geometry and flow rates will ensure more accurate DNA flow-stretching data interpretation when crowding agents are involved.

## Author Contributions

A.H. and H.K. conceived and designed this study. F.T.Z., H.Al-Z., and J.R. performed the experiments. A.H. devised the model, ran the simulations, and analyzed the data. H.K. and A.H. wrote the manuscript.

## Acknowledgements

This work was supported by the National Institutes of Health (R35GM143093 to H.K.). We acknowledge support from the National Science Foundation for the acquisition of the rheometer used in this work (DMR 2122178 to Karen Lozano).

## Supporting information

**Fig. S1.**
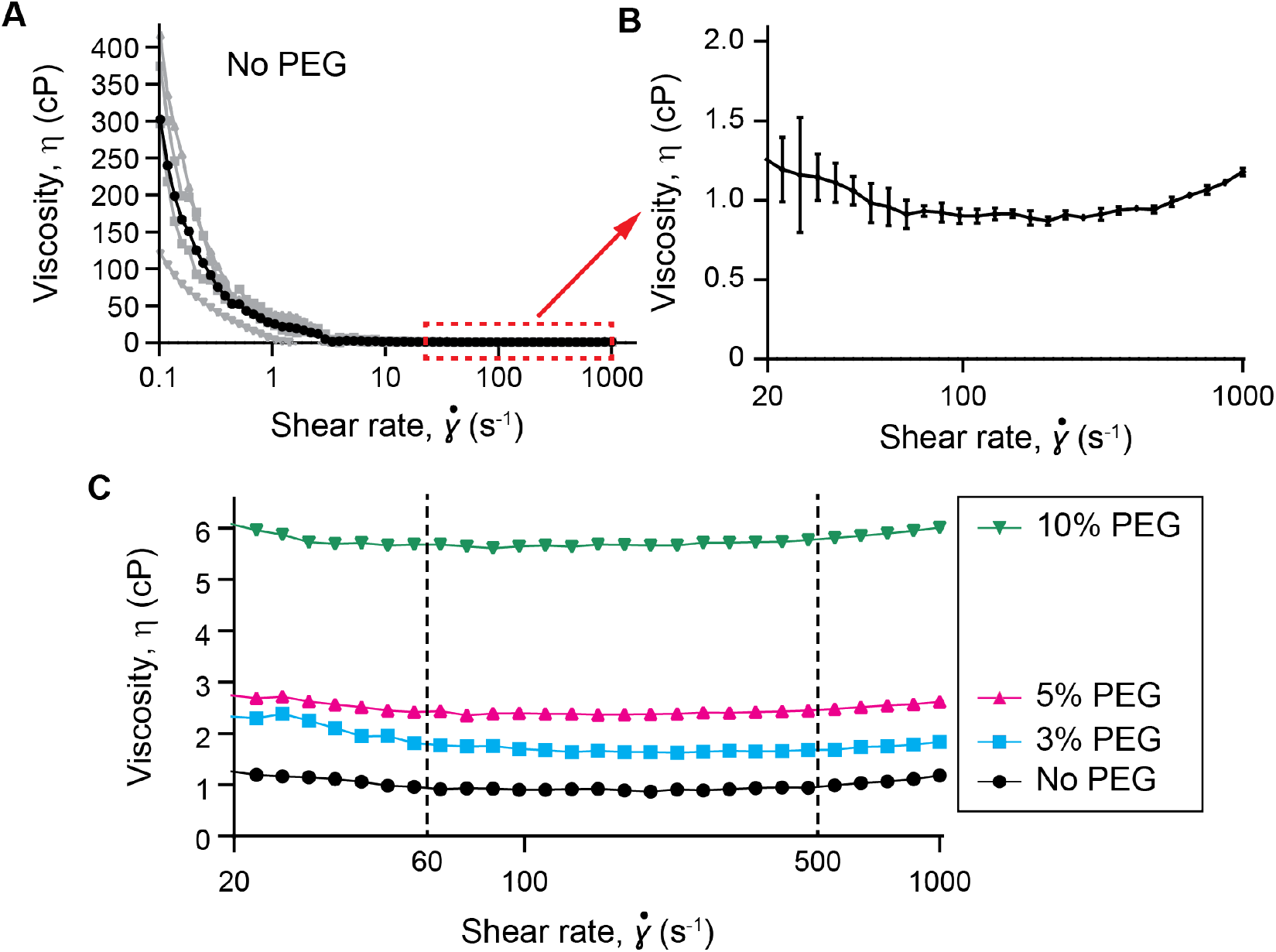
Measurements of buffer viscosity *η* as a function of shear rate 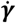 in the absence and presence of polyethylene glycol (PEG). (A) Buffer viscosity η in the absence of PEG. Shown are the results for η in units of centipoise (cP) for four measurements (gray) and the average of these measurements (black) (1 cP = 1 mP · s). (B) For better visibility, shown is the average of η in (A) for shear rates 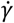 in the range 20–1000 s^-1^ as indicated by the red box in (A). Error bars: SD. (C) Buffer viscosity η in the absence of PEG and with 3%, 5%, and 10% concentrations of PEG, respectively, for shear rates 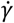 in the range 20–1000 s^-1^ obtained as the average of five measurements for each concentration of PEG. Shear rates were averaged over the range 60-500 s^-1^ for subsequent analysis in this study.

**S1 Movie. Animation of a Brownian dynamics simulation of our model chain**. The movie was obtained for a model chain with *N* = 50 beads (Fig 2) by numerical iteration of the Langevin equation (Eq. 18). Shown is the projection of the bead positions on the *xz*-plane. Initially, the chain is aligned with the *z*-axis perpendicular to the surface in the *xy*-plane (Fig 1B). It is then stretched by shear flow in the *x*-direction characterized by the Péclet number *Pe* = 0.02 (Eq. 21) corresponding to the Weissenberg number *Wi* = 333 *Pe* = 6.66 (Eqs. 22 and 23). The time steps in Eq. 18 are Δ*t* = 10^−4^ in units of *τ* _0_ (Eq. 17). The animation contains 500 frames with a delay of 10^5^ Δ*t* between frames (in units of *τ* _0_) corresponding to a total time of 5000 *τ*_0_ . Note the cyclic dynamics reported in references 18 and 20.

**S2 Movie. Flow-stretching of quantum dot-labeled bacteriophage lambda DNA**. The movie was recorded by an EMCCD camera (see main text, Section 2). The movie is 8 times faster than in real-time. The dimensions of the frame are 3.20 × 17.12 μm. In the beginning of the movie where there is no flow, the average quantum dot position corresponds to the DNA tether point location. Applying buffer stretches the surface-tethered quantum dot-labeled DNA in the direction of the buffer flow (from the bottom to the top). See Fig 1A.

